# A 37-color spectral flow cytometry panel to characterize phenotype and cytokine expression in human intestinal T cells

**DOI:** 10.1101/2025.10.29.685441

**Authors:** Ciska Lindelauf, Qinyue Jiang, Frits Koning, Andrea E. van der Meulen, Vincent van Unen, M. Fernanda Pascutti

## Abstract

We developed a 37-color spectral flow cytometry panel to characterize T cell function, with a focus CD4^+^ T helper (Th) cells, in the human intestine. The panel includes lineage-defining transcription factors, markers for memory status, tissue residency and T cell activation, and 11 cytokines representing multiple Th cell lineages. The panel is currently being used to investigate CD4^+^ Th cell function in intestinal and peripheral blood samples of inflammatory bowel disease (IBD) patients. The panel could however be applied to investigate both peripheral and tissue-resident T cell function in a range of immune-mediated diseases.

## Background

Inflammatory bowel disease (IBD), comprising Crohn’s disease (CD) and ulcerative colitis (UC), is characterized by chronic intestinal inflammation driven by dysregulated immune responses. CD4^+^ T helper (Th) cells play a central role in IBD pathogenesis, with multiple Th subsets contributing to disease in a complex and often overlapping manner, including Th1, Th2, Th9, Th17, Th22, Tfh and Tregs[1]. Understanding the balance and dysfunction of these diverse Th populations in the intestinal mucosa is critical for elucidating disease mechanisms and identifying therapeutic targets.

A major challenge in studying Th cell responses in IBD is the need to simultaneously assess both cellular phenotype and functional capacity. While tissue cytokine levels can be measured using techniques such as ELISA or multiplex assays, these approaches cannot identify which specific cell populations are producing cytokines or assess co-expression patterns of multiple cytokines at the single-cell level. Flow cytometry overcomes this limitation by enabling simultaneous detection of surface markers for phenotypic characterization alongside intracellular cytokine production. However, conventional flow cytometry panels have been constrained by spectral overlap, typically focusing either on detailed phenotyping or functional cytokine analysis, but rarely achieving comprehensive coverage of both.

The advent of spectral flow cytometry has expanded the capacity for high-dimensional single-cell analysis, allowing for the inclusion of a greater number of markers with improved resolution. Here, we present a 37-marker spectral flow cytometry panel specifically designed to comprehensively analyze circulating and intestinal CD4^+^ Th cells. This panel includes 11 cytokines representing multiple Th lineages (Th1, Th2, Th9, Th17, Th22, Tfh and regulatory responses), alongside markers for T cell memory status, tissue residency, activation status, and lineage-defining transcription factors (Tables 1 and 2). By capturing this breadth of information in a single assay, this panel enables simultaneous analysis of multiple Th subsets and their functional states, providing insights into the complex CD4^+^ T cell responses that characterize intestinal inflammation in IBD.

**Table 1.** Summary table.

|  |  |
| --- | --- |
| Purpose | Comprehensive phenotypic and functional analysis of human CD4+ Th cells |
| Species | Human |
| Cell type | Intestinal cells, PBMC, tonsil mononuclear cells |
| Cross-reference | OMIP-001, OMIP-008, OMIP-009, OMIP-014, OMIP-022, OMIP-056, OMIP-060, OMIP-067, OMIP-075, OMIP-080, OMIP-082, OMIP-091, OMIP-096, OMIP-110 |

**Table 2.** Reagent table.

| **Specificity** | **Clone** | **Fluorochrome** | **Purpose** |
| --- | --- | --- | --- |
| Viability |  | LIVE DEAD Blue | Viability |
| CD45 | HI30 | PerCP | Leukocytes |
| CD3 | UCHT1 | Pacific Blue | Pan T cell |
| CD4 | OKT4 | BV510 | CD4+ T cells |
| CD7 | M-T701 | BUV805 | T cells and ILCs |
| CD8 | RB545 | SK1 | CD8+ T cells |
| CD56 | NCAM16.2 | BUV661 | NK/NKT-like cells |
| TCR$\gamma/\delta$ | B1 | PE/Fire 810 | $\gamma/\delta$ T cells |
| TCR Vα7.2 | OF-5A12 | BV786 | MAIT cells |
| CD27 | L128 | BUV496 | T cell memory |
| CD45RA | HI100 | Spark UV 387 | T cell memory |
| CD197 (CCR7) | G043H7 | Spark NIR 685 | T cell memory |
| CD69 | FN50 | Super Bright 436 | Tissue residency and activation |
| CD103 | Ber-ACT8 | PE/Fire 700 | Tissue residency |
| CD38 | HIT2 | APC/Fire 810 | Activation |
| CD137 (4-1BB) | 4B4-1 | BV650 | Activation |
| CD154 (CD40L) | 2431 | BV605 | Activation |
| HLA-DR | G46-6 | BV480 | Activation |
| CD161 | HP-3G10 | BUV563 | T cell differentiation |
| CD294 (CRTH2) | BM16 | BUV615 | Chemokine receptor |
| FoxP3 | PCH101 | PE/Cy5 | Transcription factor |
| Helios | 22F6 | PerCP/eFluor 710 | Transcription factor |
| Ki-67 | B56 | BUV395 | Transcription factor |
| RORγt | Q21-559 | PE | Transcription factor |
| T-bet | 4B10 | RB613 | Transcription factor |
| Granzyme B | GB11 | RB744 | Cytotoxicity |
| IFNγ | B27 | BV750 | Cytokine |
| IL-2 | MQ1-17HI2 | BUV737 | Cytokine |
| IL-4 | MP4-25D2 | RB780 | Cytokine |
| IL-5 | TRFK5 | RY586 | Cytokine |
| IL-9 | MH9A3 | BV421 | Cytokine |
| IL-10 | JES3-19F1 | R718 | Cytokine |
| IL-13 | JES10-5A2 | APC | Cytokine |
| IL-17A | N49-653 | RY610 | Cytokine |
| IL-21 | 3A3-N2.1 | AF647 | Cytokine |
| IL-22 | 22URTI | PE/Cy7 | Cytokine |
| TNFα | MAb11 | Spark Blue 515 | Cytokine |

Dead cells are first excluded using LIVE/DEAD Blue, followed by selection of CD45^+^ immune cells to exclude epithelial and stromal cells commonly found in intestinal biopsies. Pan T cells are identified as CD3^+^, while this panel also distinguishes other lymphocyte subsets including ILCs (CD3^-^CD7^+^CD56^-^) and NK cells (CD3^-^CD7^+^CD56^+^) (Figure 1A-B). Given that Th cells and ILCs mirror each other in phenotype, many markers included in this panel can be applied to further characterize the ILC compartment.

**Figure 1.**
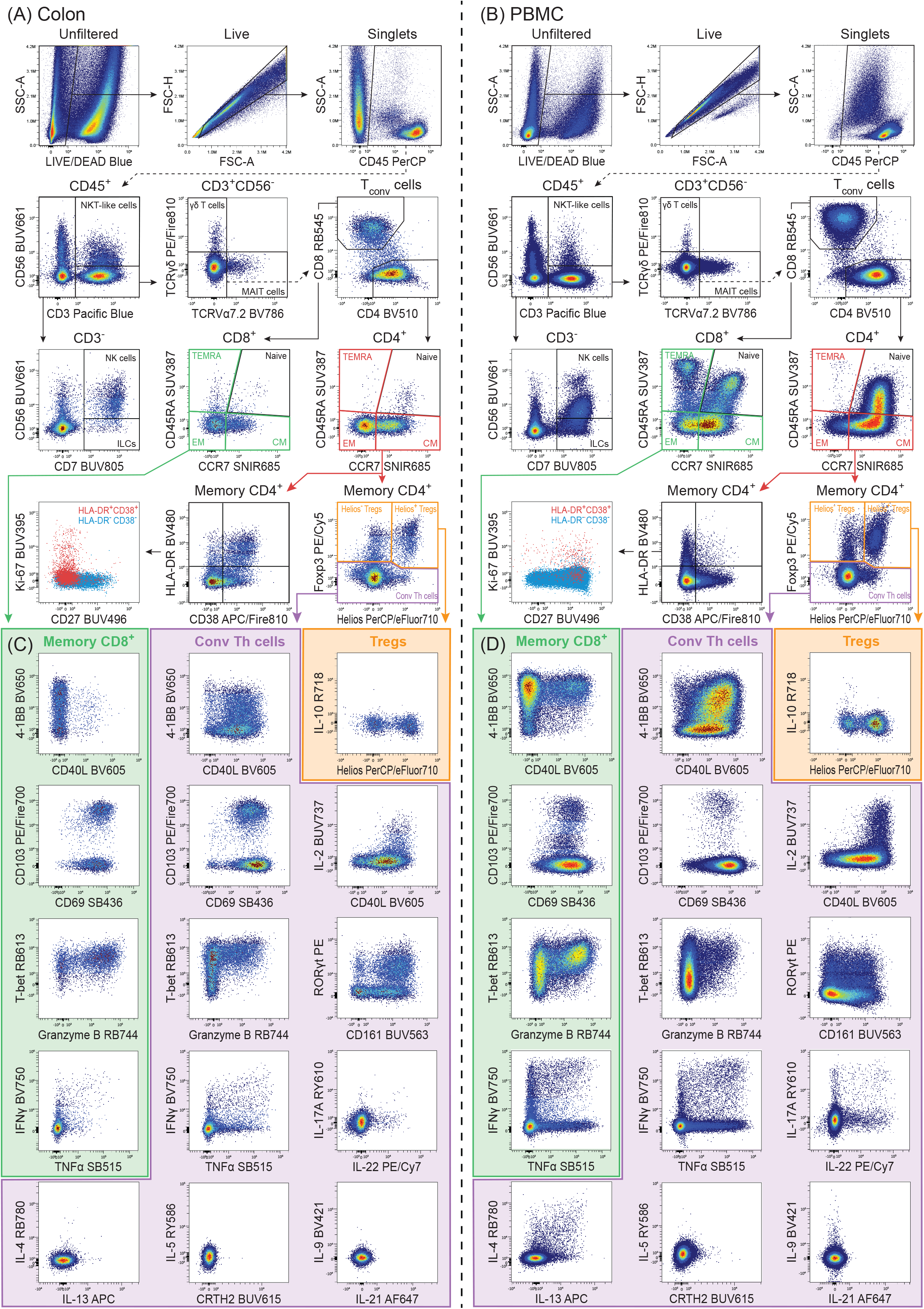
Example staining figure of 37-marker panel on human colonic cells from IBD patient with active disease (A-C) and PBMC from healthy donor (B-D). Samples were processed and stained as described in the online protocol. In short, colonic biopsies were processed into a single-cell suspension. Both colonic cells and PBMC were rested in culture overnight. The next day, cells were stimulated with anti-CD3/CD28 for 6h. Brefeldin A and BD GolgiStop were added during the final hour of stimulation, and cells were finally harvested for staining before being recorded on a 5-laser Cytek Aurora spectral flow cytometer (Cytek Biosciences). (A-B) Manual gating strategy as described in main text. (C-D) Bivariate plots of markers of activation, tissue-residency, cytokines and Th subset-defining transcription factors and surface receptors shown on (1) memory CD8^+^ T cells (2) conventional CD4^+^ Th cells and (3) CD4^+^ Tregs.

Within the T cell compartment, three main types of unconventional T cells can be identified: NKT-like (CD56^+^), γδ T (TCRγδ^+^), and MAIT (TCRVα7.2^+^) cells. Conventional T cells are divided into CD4^+^ and CD8^+^ subsets and further stratified by memory status using CD45RA and CCR7: naïve (CD45RA^+^CCR7^+^), central memory (CM, CD45RA^-^CCR7^+^), effector memory (EM, CD45RA^-^CCR7^-^), and terminally differentiated effector memory cells re-expressing CD45RA (TEMRA, CD45RA^+^CCR7^-^). The memory CD4+ compartment is further divided into Foxp3^+^ Tregs and Foxp3^-^ conventional Th cells, with Helios expression indicating stable Treg phenotype[2] (Figure 1A-B).

HLA-DR and CD38 serve as markers for T cell activation. We previously identified intestinal HLA-DR^+^CD38^+^ EM CD4^+^ T cells that are significantly enriched in IBD patients with active disease, characterized by high proliferation (Ki-67^+^) and loss of CD27 expression[3]. Similar cells have been identified in other studies and correlated with disease severity and therapy non-response[4-6]. This phenotype represents chronic immune activation and has been described in other inflammatory and infectious diseases [7-10]. The inclusion of these four markers (HLA-DR, CD38, Ki-67, and CD27) enables investigation of cytokine expression within this disease-associated T cell subset.

The costimulatory markers 4-1BB (CD137) and CD40L (CD154) are specifically and transiently upregulated upon TCR activation, allowing identification of activated T cells after in vitro stimulation. CD40L is predominantly expressed by conventional Th cells, while Tregs and CD8 T cells express 4-1BB[11-13]. CD69 and CD103 (integrin αE) are major tissue-residency markers, although CD69 is also rapidly upregulated after T cell activation. CD161, while not a tissue-residency marker per se, is upregulated on gut-resident T cells and IL-17-producing cells[14].

This panel includes 11 cytokines produced by several Th lineages: IFNγ, IL-2, IL-4, IL-5, IL-9, IL-10, IL-13, IL-17A, IL-21, IL-22, and TNFα. Additionally, Granzyme B is included to identify both CD4^+^ and CD8^+^ cytotoxic T lymphocytes (CTLs). The panel also contains several transcription factors and surface receptors to delineate Th subsets (Figure 1C-D). T-bet is upregulated in CTLs and Th1 cells, while RORγt is upregulated in Th17 cells, and surface receptor CRTH2 has been shown to reliably identify Th2 cells[15].

### Similarity to other OMIPs

OMIPs that are similar to this panel are listed in Table 1. There are several other OMIPs that have been designed for use on intestinal cells, including OMIP-082, -096 and -110[16-18]. However, these panels are not designed to detect intracellular cytokines. Other OMIPs that are designed to analyze intracellular cytokine expression in human T cells include OMIP-001, -008, -009, -014, -022, -056, -060, -067, -075, -080 and -091[19-30]. However, these panels do not contain the same breadth of T cell cytokines. In addition, what makes this panel unique is the combination of cytokine detection with markers of tissue-residency and Th subset-defining transcription factors.

## Acknowledgements

Flow cytometry was performed on 5-laser Cytek Aurora spectral analyzers (Cytek Biosciences) at the Flow cytometry Core Facility (FCF) of Leiden University Medical Center (LUMC) in Leiden, Netherlands (https://www.lumc.nl/research/facilities/fcf).

## Ethics Statement

PBMC were isolated from buffy coats collected from healthy donors via Sanquin Blood Bank (Amsterdam, Netherlands). Tonsils were obtained from routine tonsillectomies at Alrijne Hospital (Leiderdorp, Netherlands) and approved by the Medical Ethical Committee of the Leiden University Medical Center (LUMC) (protocol B21.019). Intestinal biopsies from IBD patients were obtained via the LUMC Biobank (Leiden, Netherlands) and approved by the Medical Ethical Committee of the LUMC (protocol P15.193).

## Author Contributions

**Ciska Lindelauf:** investigation (equal); methodology (equal); validation (equal); visualization (lead); writing – original draft preparation (lead). **Qinyue Jiang:** investigation (equal); methodology (equal); validation (equal); writing – review and editing (equal). **Frits Koning:** funding acquisition; supervision (supporting); writing – review & editing (supporting). **Andrea van der Meulen:** resources; supervision (supporting); writing – review & editing (supporting). **Vincent van Unen:** conceptualization (supporting); funding acquisition; project administration (equal); supervision (equal); writing – review & editing (lead). **Fernanda Pascutti:** conceptualization (lead); funding acquisition; project administration (equal); supervision (equal); writing – review & editing (supporting).

