## Supplemental materials for "A 37-color spectral flow cytometry panel to characterize phenotype and cytokine expression in human intestinal T cells"

**Panel development strategy**

This panel was developed on a 5-laser Cytek® Aurora spectral analyzer (Cytek Biosciences). The instrument configuration can be found in Online Table 1. A list of all commercially purchased detection reagents used in the final panel can be found in Online Table 2. Rather than designing a panel from scratch, we used our previously published T cell phenotyping panel as a starting point[1], which covered a broad range of transcription factors and chemokine receptors for in-depth phenotyping of freshly isolated human intestinal cells. In this new panel, many of these markers are swapped for cytokines and markers of T cell activation in order to investigate function of intestinal cells after *in vitro* stimulation. The concept was to create two panels - one with a focus on T cell phenotyping and the other on T cell function – that can be used in parallel on samples from the same patient.

During panel development, we tested several iterations of marker-fluorochrome combinations:

- CD40L was initially assigned to PE/Cy5. The positive signal was very bright, and PE/Cy5 is notorious for spillover into other channels. We therefore reassigned CD40L to a dimmer fluorochrome, BV605, which was still sufficient to separate the positive and negative population (Online Figure 1A).
- Foxp3 was initially assigned to APC. To improve the separation between the positive and negative population, we moved Foxp3 to PE/Cy5 (Online Figure 1B). This also left APC available to assign to a cytokine instead, as cytokine antibodies are often only available on a small range of fluorochromes.
- To identify Th2 cells, BUV615 was initially assigned to CCR4. It has however been reported that CRTH2 is a more reliable marker for this purpose, and some $CCR4^+$ cells can also belong to other Th subsets [2]. In a side-by-side comparison, we

observed that all CRTH2$^{+}$ CD4$^{+}$ T cells are GATA3$^{high}$, while this is not the case for all CCR4$^{+}$ cells (Online Figure 1C). We therefore swapped CCR4-BUV615 with CRTH2-BUV615.

- Th2 cytokines IL-4 and IL-13 were initially stacked together on BV421. While we were developing the panel, BD launched several new RealYellow and RealBlue fluorochromes, which allowed us to assign IL-4 and IL-13 to separate fluorochromes (Online Figure 1D) and add IL-5 on RY586. This also made BV421 available to assign to IL-9.
- TNFα was initially assigned to BUV395. We later decided to include Ki-67 in the panel to investigate T cell function within proliferating subsets. In our previous panel we assigned Ki-67 to BUV395[1]. We therefore moved TNFα to Spark Blue 515 (Online Figure 1E).
- PE was initially assigned to Tfh cell chemokine CXCL13. We tested the antibody on a range of samples (PBMC, tonsil, intestine) but were unable to identify clear expression in CD4$^{+}$ T cells. PE was therefore reassigned to RORγt (Online Figure 1F).

**Titrations**

Surface antibodies were titrated with a starting dilution 1:10 up to 1:640, with 2-fold dilution steps in between (Online Figure 2A). Intracellular markers are stained overnight, which required approximately 10-fold less antibody[3], and were therefore titrated from dilution 1:100 to 1:6400 (Online Figure 2A-C). The recommended working concentration for viability dyes is generally lower than for antibodies, and LIVE/DEAD Blue was therefore titrated from 1:500 to 1:3200 (Online Figure 2A). Due to the scarcity of available intestinal tissue, titrations were done on PBMC, or tonsillar cells for markers that were not abundantly expressed on PBMC. Cytokines and activation markers were titrated on both stimulated and unstimulated cells. Cells were rested overnight as outlined in our online protocol, and stimulated with 20 ng/mL phorbol 12-myristate 13-acetate (PMA) and 1 µg/mL ionomycin for 6h. 10 µg/mL Brefeldin A was added during the final 4h of stimulation. In some cases, cells were co-stained with a small panel of lineage identifying antibodies to asses titrations on specific cell subsets of interest, such as memory $CD4^+$ T cells. Titrations were analyzed in FlowJo™ v10.10.0 Software (BD). Stain indexes were calculated using the StainIndex v1.8.1 plugin. Titers were also assessed visually for (1) separation between the positive and negative population (2) brightness of positive population and (3) background on the negative population. Typically, antibody concentrations with the highest stain index were selected as optimal. In some cases, for primary markers, antibody concentrations with a lower stain index were selected both for economic reasons and to minimize spillover into other channels.

**Biological controls**

Intestinal samples (Figure 1A,C and Online Figure 3A,C) were stained in parallel with PBMC (Figure 1B,D and Online Figure 3B,D) and tonsil mononuclear cells (Online Figure 4A-D). In addition to cells stimulated with anti-CD3/CD28, unstimulated cells stained to assess background in cytokine channels (Online Figure 3 and 4B,D).

**Optimization of staining protocol**

In order to analyze intestinal cells with flow cytometry, tissue pieces must be digested into a single-cell suspension using enzymes, which may negatively affect staining of surface proteins due to enzymatic cleavage. We have minimized these effects by selecting enzymes that have relatively little impact on surface markers and using antibody clones that have previously been validated for labeling of enzymatically digested cells[4, 5]. In addition, we rest all samples overnight before stimulation and staining. This allows intestinal cells to recover the expression of surface markers that may have been affected by enzymatic digestion[6]. It is also recommended for cryopreserved cells to recover their functional activity after thawing [7, 8].

Upon T cell activation, the TCR complex is rapidly internalized and later degraded in lysosomes [9]. CD3, TCRVα7.2 and TCRγδ are therefore not detectable on the surface of recently activated T cells. This problem can be circumvented by staining these targets intracellularly (Online Figure 5). Though TCRγδ clone 11F2 is reportedly the superior clone for detection of γδ T cells [10], we observed a decrease in staining quality when it is used after fixation/permeabilization. Subsequently, we switched to clone B1 which performed better for intracellular staining (Online Figure 5). In addition, we observed downregulation of CD4 – though not for CD8 – on activated T cells and found that staining this marker intracellularly improved staining quality (Online Figure 5).

There is a variety of commercially available fixation/permeabilization buffer kits that are recommended either for staining of nuclear or cytoplasmic proteins. We compared the performance of our panel when using eBioscience™ Foxp3 / Transcription Factor Staining Buffer Set (Invitrogen) or BD Cytofix/Cytoperm™ Fixation/Permeabilization Kit (BD Biosciences), which are recommended for staining nuclear or cytoplasmic proteins respectively. Though using the cytoplasmic protein staining kit did slightly improve the detection of some cytokines, the staining quality of transcription factors was insufficient (Online Figure 6). As our panel includes both transcription factors and cytokines, we decided to use the eBioscience™ Foxp3 / Transcription Factor Staining Buffer Set.

**Optimization of stimulation protocol**

There are several variables to consider for cytokine detection in activated T cells, such as stimulant selection, stimulation time and the use of protein transport inhibitors (PTI). We compared three stimulants—αCD3(clone OKT3)/αCD28, αCD3(clone HIT3a)/αCD28, and phorbol 12-myristate 13-acetate (PMA)/ionomycin—across two time points: 6 and 24 hours. Stimulation with PMA/ionomycin resulted in significant cell death and more severe downregulation of CD4, which hindered the identification of $CD4^+$ T cells (Online Figure 7A-B). Stimulating cells for 24h did not improve detection of most cytokines—except for IL-9 and IL-10—compared to 6h (Online Figure 7C). We found αCD3 clone HIT3a to perform slightly better than clone OKT3 (Online Figure 7C). Finally, we chose stimulation with αCD3/αCD28 (clone HIT3a) for 6h as the optimal method for our panel.

During stimulation, a protein transport inhibitor (PTI) must be added to block cytokine secretion. Two widely used PTIs are brefeldin A and monensin, which both target different steps in intracellular protein transport[11]. We stimulated PBMC with aCD3/aCD28 for 6h, adding brefeldin A and/or monensin at different timepoints during stimulation, and measured the expression of co-stimulatory molecules, cytokines and granzyme. Adding PTIs only during the final hour of stimulation increased the detection of 4-1BB, IL-4, IL-5, IL-13 and IL-22, compared to adding PTIs for the full six hours of stimulation (Online Figure 8A). Delaying the addition of  PTIs allows for auto- and paracrine IL-2 signaling which promotes cytokine production[12]. Notable is that we also observed an effect of PTIs on the median fluorescence intensity (MFI) of CD3, TCRVα7.2 and TCRγδ. In this case, the MFI was higher when monensin was added, alone or in combination with Brefeldin A, during stimulation. Delayed addition of PTIs resulted in a decrease in MFI (Online Figure 8B). In addition to blocking cytokine secretion, monensin also deacidifies endosomal and lysosomal compartments[13], and may therefore prevent degradation of internalized TCR complexes. For our final protocol, we decided to use a combination of brefeldin A and monensin added during the final hour of stimulation.

### Methodology

MATERIALS

- HBSS (Gibco catalog #14175-053)
- EDTA (0.5 M), pH 8.0, RNase-free (Invitrogen catalog # AM9260G)
- RPMI 1640 (Gibco catalog #22409-015)
- L-Glutamine, 200 mM (Gibco catalog #25030-024)
- Fetal Calf Serum (FCS) (Bodinco catalog #S00FD10003)
- IMDM (Capricorn catalog #IMDM-A)
- Collagenase Type 4 (Worthington catalog #LS004189)
- DNAse I (Roche catalog #10104159001)
- 70 µm Cell Strainer (Falcon catalog #352350)
- DPBS (Gibco catalog #14190-136)
- Acridine Orange/Propidium Iodide Stain (Aligned Genetics catalog #F23001)
- Human Serum (Sigma-Aldrich catalog #H5667-20ML)
- Penicillin-Streptomycin 10,000 U/mL (Gibco catalog #15140122)
- Normocin (Invivogen catalog #ant-nr-1)
- AffiniPure® Goat Anti-Mouse IgG, Fcγ fragment specific (Jackson ImmunoResearch, cat #115-005-071)

- CD3 Monoclonal Antibody (HIT3a), Functional Grade (eBioscience catalog #16-0039-85)
- CD28 Monoclonal Antibody (CD28.2), Functional Grade (eBioscience catalog #16-0289-85)
- Brefeldin A (Sigma-Aldrich catalog #B5936-200UL)
- BD GolgiStop™ Protein Transport Inhibitor (BD catalog #554724)
- PBS (Fresenius Kabi catalog #M090001/03 NL)
- Human TruStain FcX™ (Biolegend catalog #422302)
- BD Horizon™ Brilliant Stain Buffer Plus (BD catalog #566385)
- CellBlox™ Plus Blocking Buffer (Invitrogen catalog #C001T06F01)
- eBioscience™ Foxp3 / Transcription Factor Staining Buffer Set (Invitrogen catalog #00-5523-00)
- Falcon® 5 mL Round Bottom Polystyrene Test Tube, with Cell Strainer Snap Cap (Corning catalog #352235)

PREPARED REAGENTS

- Complete RPMI: add 5 mL L-Glutamine (200 mM) to 500 mL of RPMI 1640
- DNAse I solution: add 10 mL ultrapure water to 100 mg DNAse I to create a 10 mg/mL stock solution. Vortex until dissolved, then sterilize solution through a 0.20 µm syringe filter.
- Collagenase Type 4 solution: add 5 mL HBSS to 5000 U Collagenase Type 4 to create a 1000 U/mL stock solution. Vortex until dissolved, then sterilize solution through a 0.20 µm syringe filter.

- Digestion Mix 1: add 50 µL EDTA solution (0.5 M) to 5 mL HBSS to create a working solution of 5 mM EDTA
- Digestion Mix 2: to 5 mL IMDM, add 250 µL of Collagenase Type 4 solution (1000 U/mL stock, 50 U/mL working conc.) and 100 µL of DNAse I solution (10 mg/mL stock, 200 µg/mL working conc.)
- Culture Medium: to 45 mL IMDM, add 5 mL human serum (10% working conc.), 500 µL Pen Strep (10,000 U/mL stock, 100 U/mL working conc.), 100 µL Normocin (50 mg/ml, 100 µg/mL working conc.) and 1 mL DNAse I solution (10 mg/mL stock, 200 µg/mL working conc.)
- Staining buffer: to 50 mL PBS, add 500 µL FCS (1% working conc.), 50 µL Brefeldin A (10 mg/mL stock, 10 µg/mL working conc.) and 33 µL BD GolgiStop™ (1:1500)
- Foxp3 Fixation/Permeabilization working solution: add 1 part Foxp3 Fixation/Permeabilization Concentrate to 3 parts Foxp3 Fixation/Permeabilization Diluent
- Permeabilization Buffer: add 1 part 10X Permeabilization Buffer to 9 parts ultrapure water

DAY 1 - PROCESSING INTESTINAL BIOPSIES INTO SINGLE-CELL SUSPENSIONS

**To acquire enough cells for stimulation, collect at least four endoscopic biopsies per intestinal region. Keep biopsies separated according to their region and whether the region was visibly inflamed or uninflamed.**

1. For each region: collect at least four endoscopic biopsies in a 5 mL tube containing HBSS+1%FCS
2. Centrifuge at 300 x *g* for 8 min

3. Aspirate supernatant and add 1 mL of Digestion Mix 1 to biopsies
4. Incubate on a roller, at 37°C, for 30 min
5. Vortex biopsies for 10 sec
6. Add 4 mL IMDM to biopsies
7. Centrifuge at 300 x *g* for 8 min
8. Aspirate supernatant and add 1 mL of Digestion Mix 2 to biopsies
9. Incubate on a roller, at 37°C, for 1h
    a. Thaw reference samples during incubation time. See protocol below.
10. Vortex biopsies for 10 sec
11. Add 1 mL IMDM+20%FCS to biopsies
12. Pass biopsies and supernatant through 70 µm cell strainer into a 50 mL tube. Mash the biopsies with a syringe plunger, alternate with rinsing the cell strainer with up to 13 mL IMDM.
13. After mashing, transfer cell suspension from 50 mL to a 15 mL tube
14. Centrifuge at 300 x *g* for 8 min
15. Aspirate supernatant and resuspend pellet in 1 mL culture medium
16. Count cells in Acridine Orange/Propidium Iodide Stain on a LUNA-FX7™ Automated Cell Counter (Aligned Genetics) according to manufacturer's instructions
17. Dilute cell suspension with culture medium to a final concentration of 1-2x$10^6$ cells/mL
18. Seed cells in a TC-treated 24-well plate, 1 mL/well
19. Incubate, at 37°C, overnight
20. The next day, continue with stimulation protocol

DAY 1 - THAWING FROZEN CELLS (PBMC and tonsil)

1. Per cryovial, prepare one 15 mL tube containing 1 mL complete RPMI and 1 mL FCS
2. Thaw the cryovial in a 37°C water bath until only a small piece of ice remains
3. Transfer the contents of the cryovial (1 mL) to the prepared 15 mL tube
4. Dropwise, add 12 mL complete RPMI to the 15 mL tube
5. Centrifuge at 300 x *g* for 8 min
6. Discard supernatant and resuspend cells in 1 mL culture medium
7. Count cells in Acridine Orange/Propidium Iodide Stain on a LUNA-FX7™ Automated Cell Counter (Aligned Genetics) according to manufacturer's instructions
8. Dilute cell suspension with culture medium to a final concentration of 1-2x$10^6$ cells/mL
9. Seed cells in a TC-treated 24-well plate, 1 mL/well
10. Incubate, at 37°C, overnight
11. The next day, continue with stimulation protocol

DAY 1 – PLATE COATING

1. To a non-treated 24-well plate, add 1 mL/well of PBS supplemented with 4 µg/mL anti-mouse IgG
2. Incubate, at 37°C, for 2h
3. Wash coated wells twice:
    a. Aspirate supernatant
    b. Add 1 mL PBS
    c. Repeat steps a-b
4. Aspirate supernatant and add 1 mL/well of PBS+10%FCS
5. Incubate, at 37°C, for 30 min
6. Aspirate supernatant and add 1 mL/well of PBS supplemented with 2.5 µg/mL anti-CD3

7. Incubate, at 4°C, overnight

DAY 2 - STIMULATION

8. Aspirate supernatant from anti-CD3 coated wells
9. Transfer 1 mL of cells from the TC-treated plate to the anti-CD3 coated plate, do this for each sample type
    a. For each sample type, you should have at least 1 well with cells in the TC-treated plate that serves as an unstimulated control, and 1 well with cells in the anti-CD3 coated plate for stimulation.
10. Add 1 µL anti-CD28 (1 mg/mL stock, 1 µg/mL working conc.) to each anti-CD3 coated well
11. Incubate, at 37°C, for 5h
12. Add 1 µL Brefeldin A (10 mg/mL stock, 10 µg/mL working conc.) and 0.67 µL BD GolgiStop™ (1:1500) to all wells, both with and without anti-CD3 coating
13. Incubate, at 37°C, for 1h
14. Transfer cells from 24-well plates to 1.5 mL tubes
15. Rinse each well with 0.5 mL HBSS and add this to the 1.5 mL tubes
16. Centrifuge at 300 x *g* for 8 min
17. Aspirate supernatant and resuspend pellets in 1 mL HBSS
    a. From each sample tube, transfer 100 µL to a new 1.5 mL tube. This aliquot will serve as the unstained (US) sample. The remaining 900 µL is the multicolor (MC) sample.
18. Centrifuge at 300 x *g* for 8 min
19. Aspirate supernatant and continue with staining protocol

DAY 2 - STAINING

1. Resuspend each MC sample in 20 µL of HBSS with LIVE DEAD Blue (1:500, see Online Table 2)
    a. Resuspend US samples in 20 µL staining buffer.
2. Incubate protected from light, at RT, for 10 min
3. Add 1 µL Human TruStain FcX™ to each MC sample
4. Incubate protected from light, at RT, for 10 min
5. Prepare the surface antibody mix with 10 µL Brilliant Stain Buffer Plus and 5 µL CellBlox™ Plus Blocking Buffer for each MC tube, add respective surface antibodies calculated with a final volume of 100 µL per MC tube (see Online Table 2). Add staining buffer to the antibody mix to bring the volume to 80 µL calculated for each MC tube.
6. Add 80 µL surface antibody mix to each MC tube; this brings the final staining volume to 100 µL
    a. Add 80 µL staining buffer to each US tube
7. Incubate protected from light, at RT, for 1h
8. Wash samples twice:
    a. Add 200 µL staining buffer to each tube
    b. Centrifuge 300 x *g* for 5 min
    c. Discard supernatant and resuspend samples in 200 µL staining buffer
    d. Centrifuge 300 x *g* for 5 min
    e. Discard supernatant
9. Resuspend each sample in 100 µL Foxp3 Fixation/Permeabilization working solution
10. Incubate protected from light, at 4℃, for 30 min
11. Wash samples twice:

   a. Add 200 µL Permeabilization Buffer to each tube
   b. Centrifuge 700 x *g* for 8 min
   c. Discard supernatant and resuspend samples in 200 µL Permeabilization Buffer
   d. Centrifuge 700 x *g* for 8 min
   e. Discard supernatant
12. Prepare the surface antibody mix with 10 µL Brilliant Stain Buffer Plus and 5 µL CellBlox™ Plus Blocking Buffer for each MC tube, add respective intracellular antibodies calculated with a final volume of 100 µL per MC tube (see Online Table 2). Add Permeabilization Buffer to the antibody mix to bring the volume to 100 µL calculated for each MC tube.
13. Resuspend each MC sample in 100 µL intracellular antibody mix
   a. Resuspend each US sample in 100 µL Permeabilization Buffer
14. Incubate protected from light, at 4°C, overnight

DAY 3 – WASH AND ACQUIRE

15. Wash samples twice:
   a. Add 200 µL Permeabilization Buffer to each tube
   b. Centrifuge 700 x *g* for 8 min
   c. Discard supernatant and resuspend samples in 200 µL Permeabilization Buffer
   d. Centrifuge 700 x *g* for 8 min
   e. Discard supernatant
16. Resuspend each MC sample in 400 µL staining buffer and each US sample in 200 µL staining buffer

17. Filter each sample by pipetting it into a 5 mL round-bottom tube with 35 μm cell strainer cap
18. Store samples protected from light, on ice, and acquire the same day on a 5-laser Cytek Aurora (Cytek Biosciences)

SINGLE-STAINS

Viability single-stains were recorded on CellTraX Tox (Slingshot Biosciences) cell mimics. SpectraComp® XT (Slingshot Biosciences) hydrogel particles were used for antibody single-stains. In some cases, especially for very bright markers, single-stains were recorded on PBMC as unmixing performed better than with hydrogel particles (CD45RA - Spark UV 387, CD7 - BUV805, CD69 – Super Bright 436, HLA-DR - BV480, CD40L - BV605, TNFα – Spark Blue 515, CD8 - RB545, Granzyme B - RB744, RORγt - PE, Foxp3 - PE/Cy5, CD103 - PE/Fire 700, CCR7 - Spark NIR 685). Single-stain samples were stained using the same protocol as for multicolor samples. To ensure that single-stains were bright enough, an antibody dilution of 1:10 was used. When staining cell mimics and hydrogel particles, Human TruStain FcX™, Brilliant Stain Buffer Plus and CellBlox™ Plus Blocking Buffer were omitted from the protocol.

SOFTWARE AND ANALYSIS

Raw data was unmixed using SpectroFlo® software (Cytek Biosciences) with autofluorescence extraction. Unmixed data was analyzed in OMIQ (Dotmatics). Before formal analysis, data was cleaned using PeacoQC[14]. This was applied on compensated and scaled data, and events with marginal values were removed.

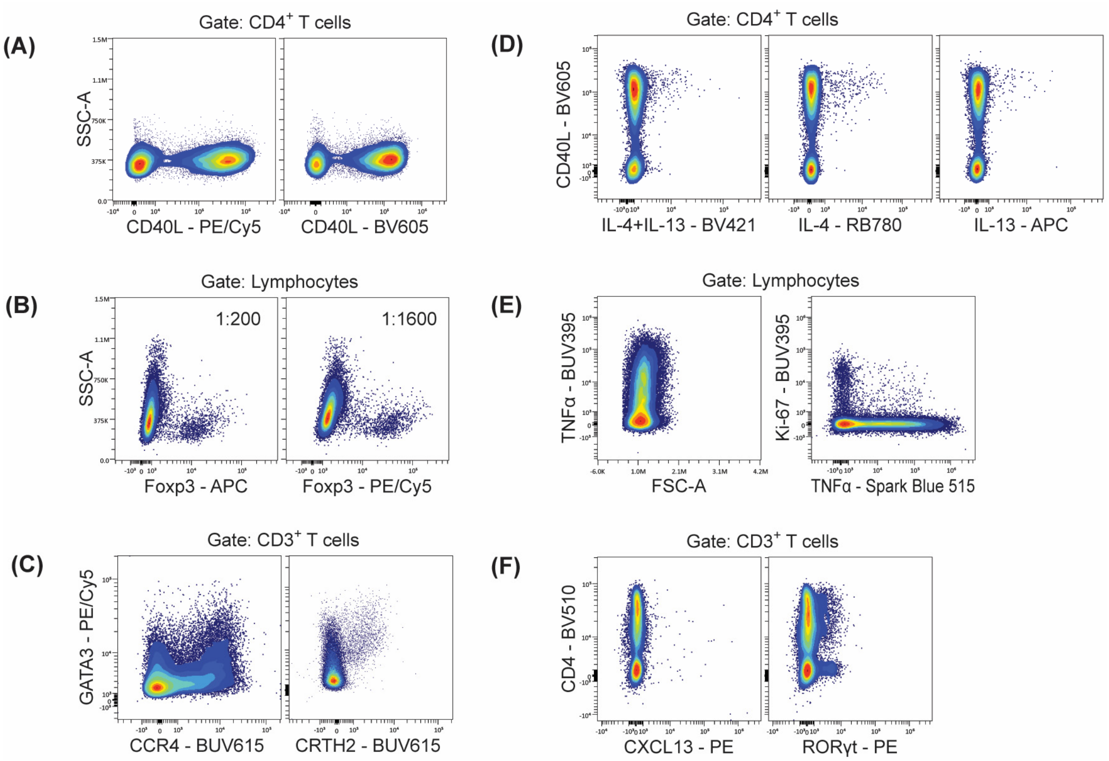


**Online Figure 1: Panel iterations.** (**A**) CD40L on PE/Cy5 was considered too bright and therefore moved to BV605. (**B**) Foxp3 was moved from APC to PE/Cy5 to increase resolution. (**C**) To identify Th2 (GATA3$^{high}$) cells, CCR4 – BUV615 was replaced with CRTH2 – BUV615. (**D**) IL-4 and IL-13 were initially stacked on the same fluorochrome, BV421, but were later separated onto RB780 and APC respectively. (**E**) TNFα originally occupied BUV395, but when Ki-67 – BUV395 was later added to the panel, TNFα was moved to Spark Blue 515. (**F**) PE was originally occupied by CXCL13, but due to lack of clear positive signal it was replaced with RORγt. Marker-fluorochrome combinations were tested on unstimulated PBMC (**B-C**), or on PBMC (**A, D, F**) and tonsil mononuclear cells (**E**) that were stimulated with PMA/ionomycin for 6h, including Brefeldin A for the final 4h of stimulation.

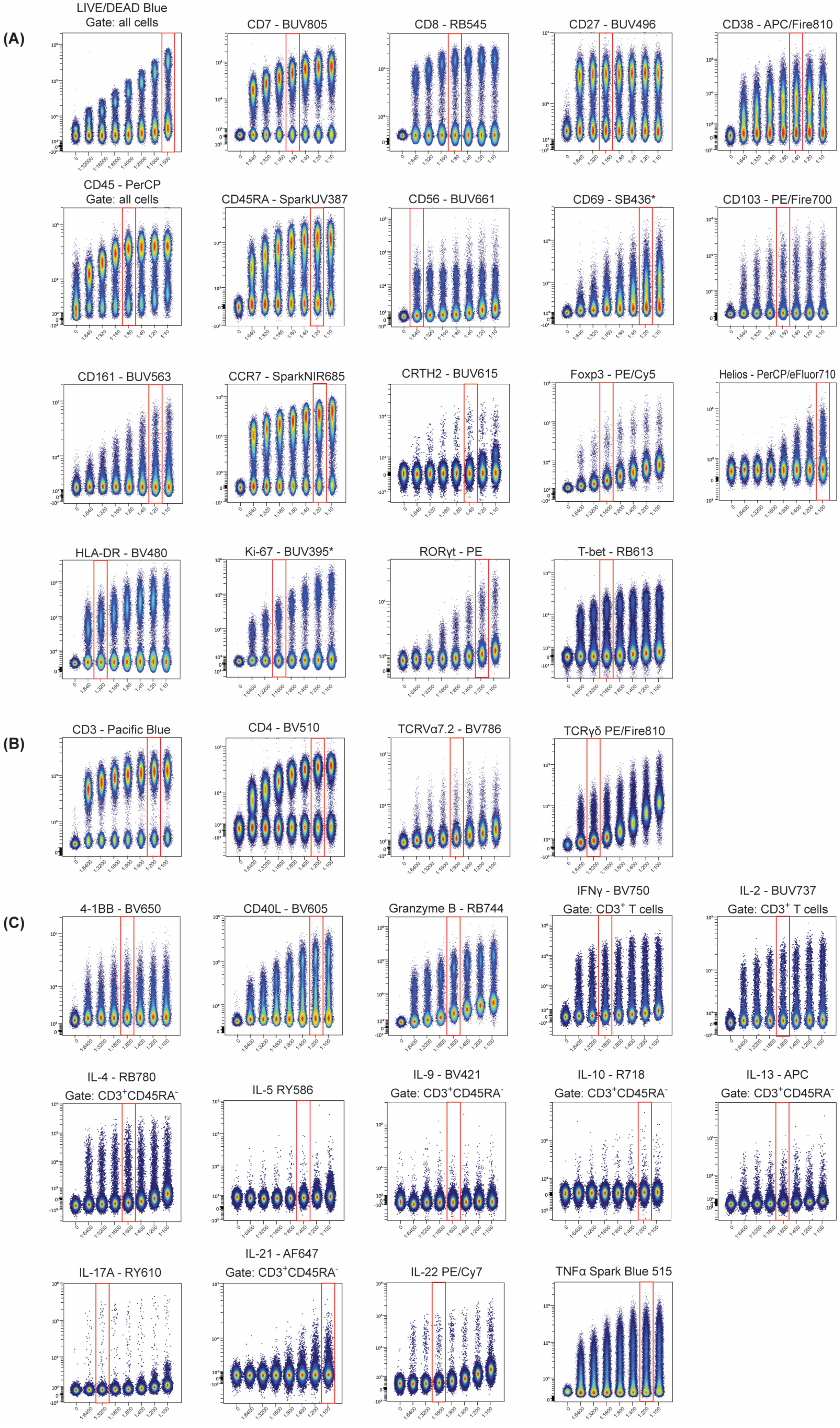
(A)
LIVE/DEAD Blue
Gate: all cells
CD7 - BUV805
CD8 - RB545
CD27 - BUV496
CD38 - APC/Fire810
CD45 - PerCP
Gate: all cells
CD45RA - SparkUV387
CD56 - BUV661
CD69 - SB436*
CD103 - PE/Fire700
CD161 - BUV563
CCR7 - SparkNIR685
CRTH2 - BUV615
Foxp3 - PE/Cy5
Helios - PerCP/eFluor710
HLA-DR - BV480
Ki-67 - BUV395*
RORγt - PE
T-bet - RB613
(B)
CD3 - Pacific Blue
CD4 - BV510
TCRVα7.2 - BV786
TCRγδ PE/Fire810
(C)
4-1BB - BV650
CD40L - BV605
Granzyme B - RB744
IFNγ - BV750
Gate: CD3+ T cells
IL-2 - BUV737
Gate: CD3+ T cells
IL-4 - RB780
Gate: CD3+CD45RA-
IL-5 RY586
IL-9 - BV421
Gate: CD3+CD45RA-
IL-10 - R718
Gate: CD3+CD45RA-
IL-13 - APC
Gate: CD3+CD45RA-
IL-17A - RY610
IL-21 - AF647
Gate: CD3+CD45RA-
IL-22 PE/Cy7
TNFα Spark Blue 515

**Online Figure 2: Titrations**. Reagents were titrated on thawed human PBMC or *tonsil mononuclear cells. Titrations followed the same staining protocol as described for multicolor samples. In some cases, samples were co-stained with a small panel of lineage-identifying markers, such as LIVE/DEAD, CD3, CD4, CD45RA and CD40L, in order to assess titrations on specific cell subsets. Titrations are shown on lymphocyte gate, unless stated otherwise. Antibodies in figure (**A**) were titrated on unstimulated cells; in (**B**) cells were stimulated with anti-CD3/CD28 and in (**C**) with PMA/ionomycin, for 6h. Brefeldin A was added during the final 4h of stimulation.

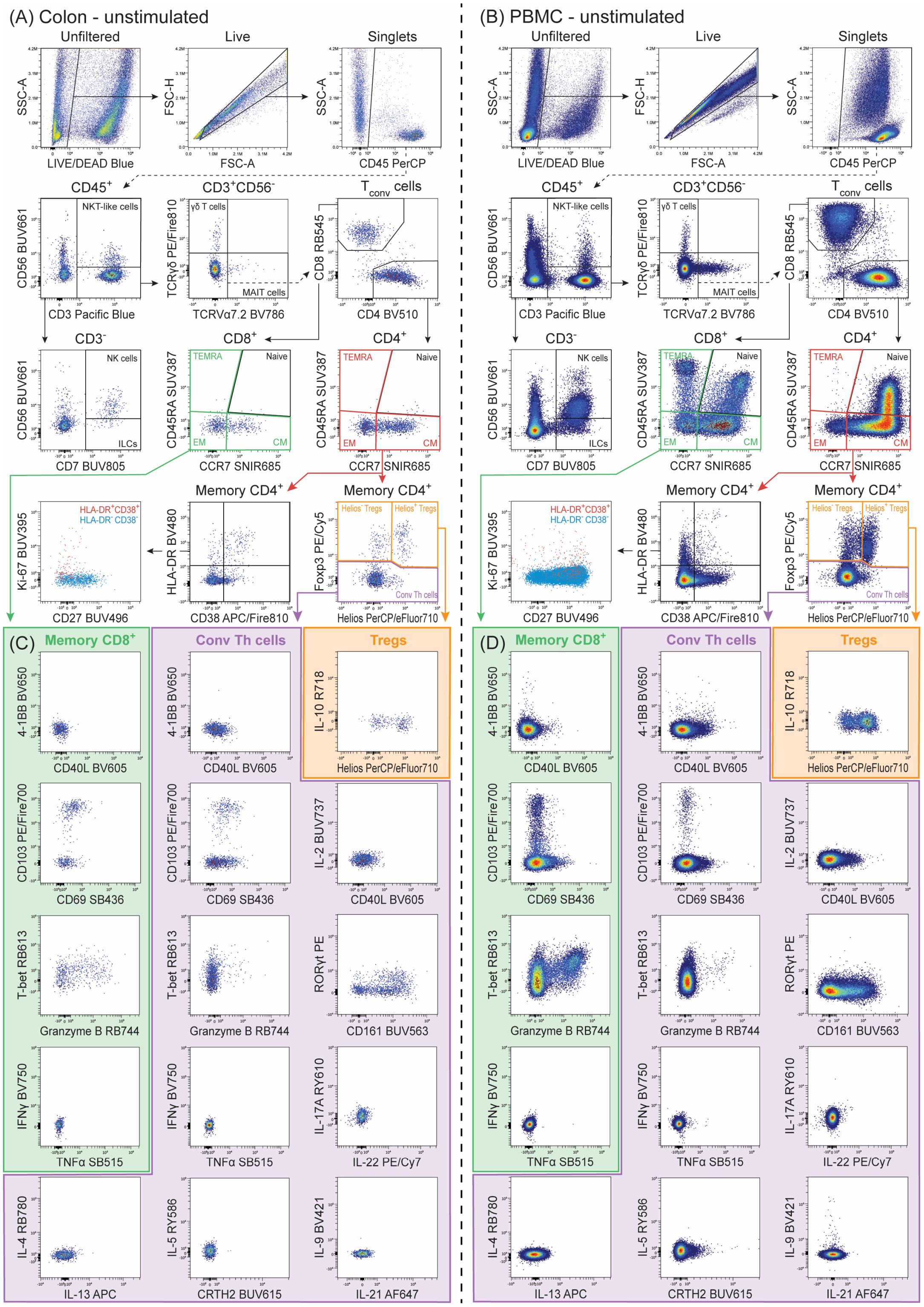
(A) Colon - unstimulated
(B) PBMC - unstimulated
Unfiltered
Live
Singlets
SSC-A
FSC-H
LIVE/DEAD Blue
FSC-A
CD45 PerCP
CD45+
CD3+CD56-
Tconv cells
CD56 BUV661
NKT-like cells
CD3 Pacific Blue
TCRγδ PE/Fire810
γδ T cells
MAIT cells
TCRVα7.2 BV786
CD8 RB545
CD4 BV510
CD3-
CD8+
CD4+
NK cells
ILCs
CD7 BUV805
CD45RA SUV387
TEMRA
Naive
EM
CM
CCR7 SNIR685
Memory CD4+
Ki-67 BUV395
HLA-DR+CD38+
HLA-DR-CD38-
CD27 BUV496
HLA-DR BV480
CD38 APC/Fire810
Foxp3 PE/Cy5
Helios- Tregs
Helios+ Tregs
Conv Th cells
Helios PerCP/eFluor710
(C)
(D)
Memory CD8+
Conv Th cells
Tregs
4-1BB BV650
CD40L BV605
IL-10 R718
CD103 PE/Fire700
CD69 SB436
IL-2 BUV737
T-bet RB613
Granzyme B RB744
RORγt PE
CD161 BUV563
IFNγ BV750
TNFα SB515
IL-17A RY610
IL-22 PE/Cy7
IL-4 RB780
IL-13 APC
IL-5 RY586
CRTH2 BUV615
IL-9 BV421
IL-21 AF647

**Online Figure 3.** Example staining figure on unstimulated human colonic cells from IBD patient with active disease (A-C) and PBMC from healthy donor (B-D). (A-B) Manual gating strategy as described in main text. (C-D) Bivariate plots of markers of activation, tissue-residency, cytokines and Th subset-defining transcription factors and surface receptors shown on (1) memory $CD8^+$ T cells (2) conventional $CD4^+$ Th cells and (3) $CD4^+$ Tregs.

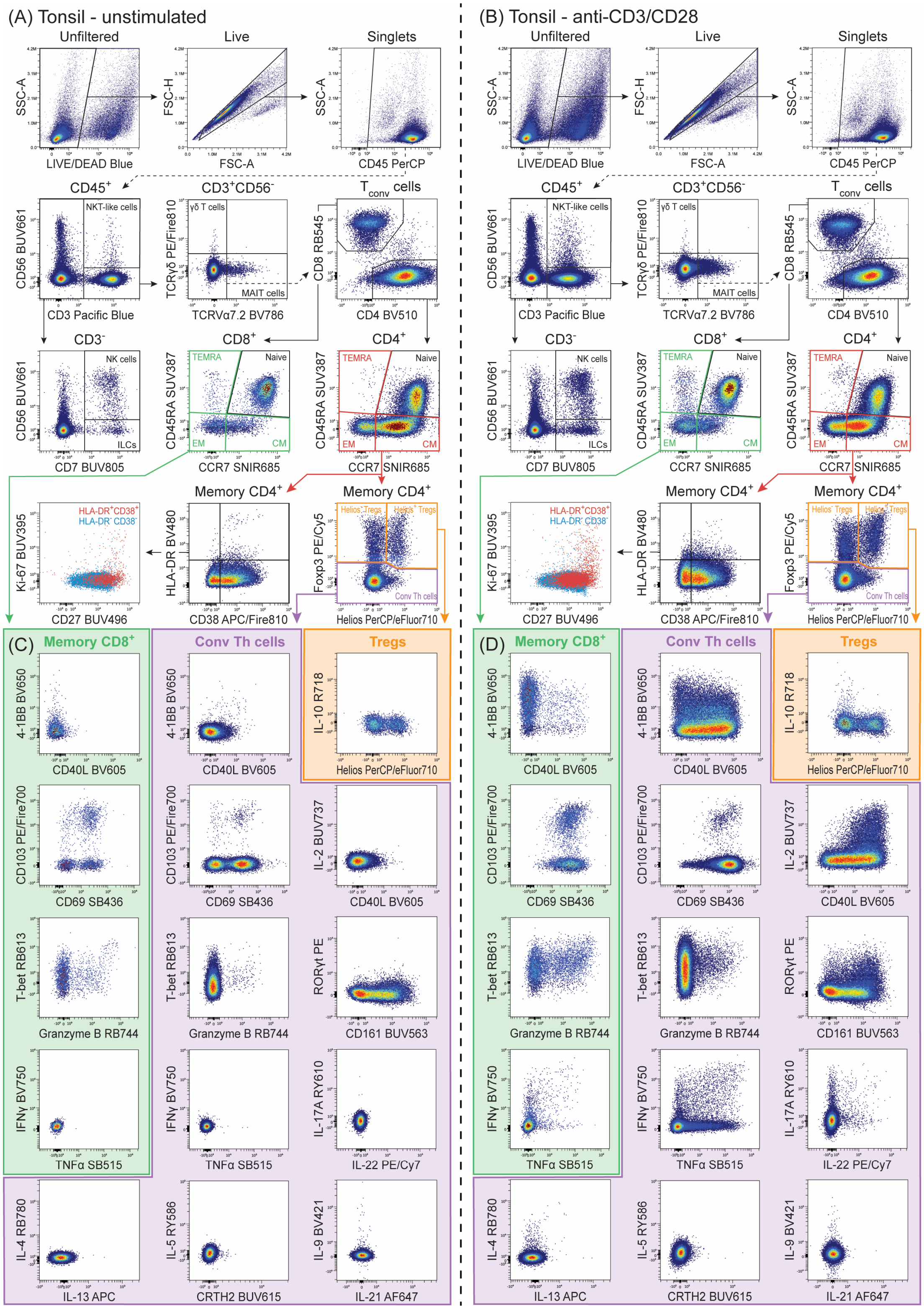

(A) Tonsil - unstimulated
Unfiltered
Live
Singlets
SSC-A
LIVE/DEAD Blue
FSC-H
FSC-A
CD45 PerCP
CD45+
CD3+CD56-
Tconv cells
NKT-like cells
CD56 BUV661
CD3 Pacific Blue
γδ T cells
TCRγδ PE/Fire810
MAIT cells
TCRVα7.2 BV786
CD8 RB545
CD4 BV510
CD3-
CD8+
CD4+
NK cells
ILCs
CD7 BUV805
TEMRA
Naive
EM
CM
CD45RA SUV387
CCR7 SNIR685
Memory CD4+
HLA-DR+CD38+
HLA-DR-CD38-
Ki-67 BUV395
CD27 BUV496
HLA-DR BV480
CD38 APC/Fire810
Helios- Tregs
Helios+ Tregs
Conv Th cells
Foxp3 PE/Cy5
Helios PerCP/eFluor710
(B) Tonsil - anti-CD3/CD28
(C) Memory CD8+
Conv Th cells
Tregs
4-1BB BV650
CD40L BV605
IL-10 R718
CD103 PE/Fire700
CD69 SB436
IL-2 BUV737
T-bet RB613
Granzyme B RB744
RORγt PE
CD161 BUV563
IFNγ BV750
TNFα SB515
IL-17A RY610
IL-22 PE/Cy7
IL-4 RB780
IL-13 APC
IL-5 RY586
CRTH2 BUV615
IL-9 BV421
IL-21 AF647
(D) Memory CD8+

**Online Figure 4.** Example staining figure on human tonsil mononuclear cells, unstimulated (A-C) and anti-CD3/CD28 stimulated (B-D). (A-B) Manual gating strategy as described in main text. (C-D) Bivariate plots of markers of activation, tissue-residency, cytokines and Th subset-defining transcription factors and surface receptors shown on (1) memory $CD8^+$ T cells (2) conventional $CD4^+$ Th cells and (3) $CD4^+$ Tregs.

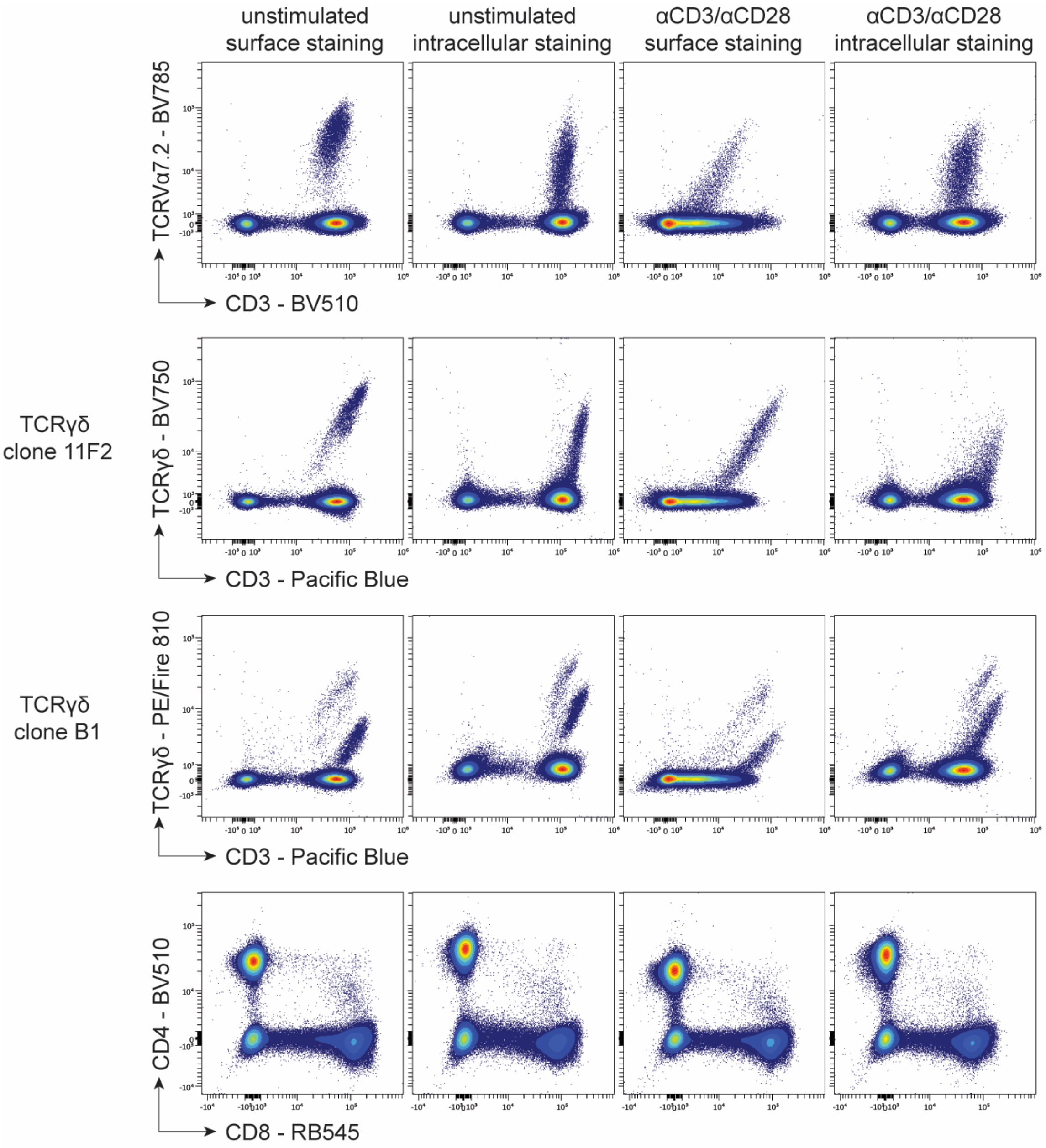


**Online Figure 5: Staining optimization of TCRs and co-receptors.** Surface and intracellular staining of TCRVα7.2, TCRγδ and co-receptors CD3, CD4 and CD8 were compared on PBMC, both unstimulated and stimulated with anti-CD3/CD28 for 6h. Brefeldin A and BD GolgiStop™ (containing monensin) were added during the final hour of stimulation.

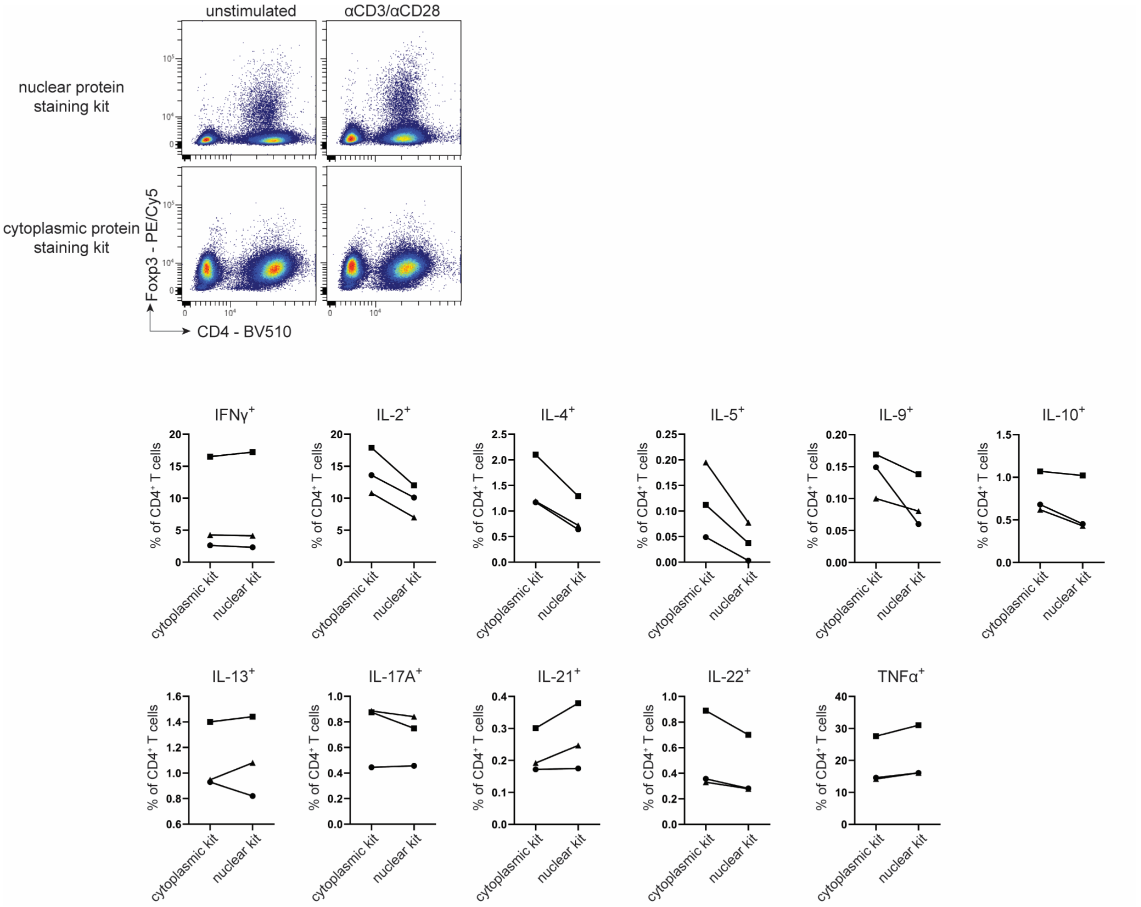


**Online Figure 6: Comparison of intracellular staining kits.** PBMC were stimulated with PMA/ionomycin for 6h and Brefeldin A was included during the final 4h of stimulation. Before intracellular staining, cells were fixed and permeabilized either with eBioscience™ Foxp3 / Transcription Factor Staining Buffer Set (Invitrogen) or BD Cytofix/Cytoperm™ Fixation/Permeabilization Kit (BD Biosciences), which are recommended for staining nuclear or cytoplasmic proteins, respectively. Data from 3 healthy donors.

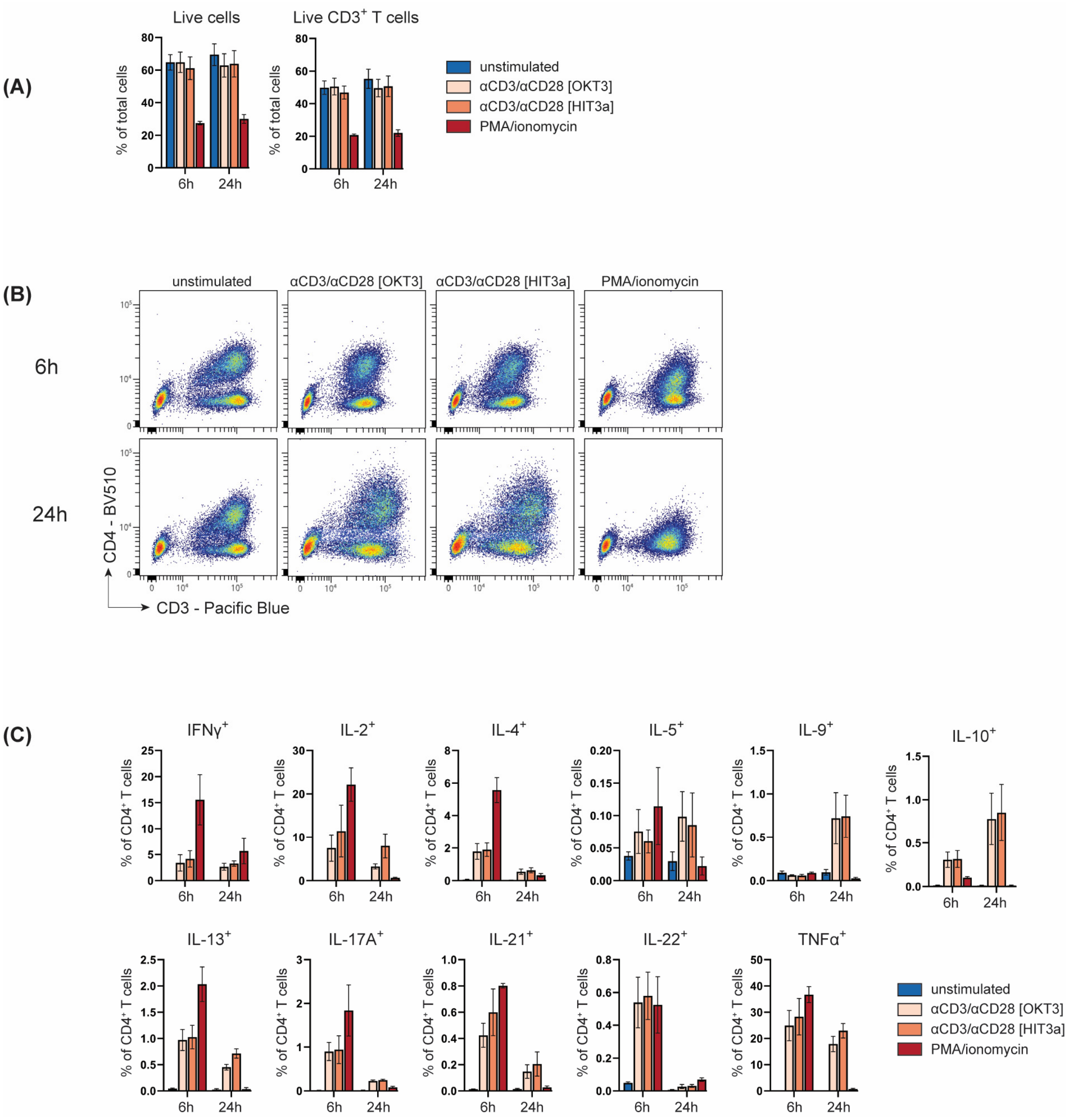


**Online Figure 7: Comparison of stimulation methods and duration.** Three stimulants—αCD3(clone OKT3)/αCD28, αCD3(clone HIT3a)/αCD28, and PMA/ionomycin—across two time points—6 and 24 hours—were compared on PBMC. For each condition, Brefeldin A was added during the final 4h of stimulation. (**A**) Yield of total live cells and $CD3^+$ T cells is shown for each condition. (**B**) Representative plots of CD4 signal, shown on live lymphocyte gate. (**C**) Quantification of intracellular cytokines within the $CD4^+$ T cell compartment. Data from 3 healthy donors.

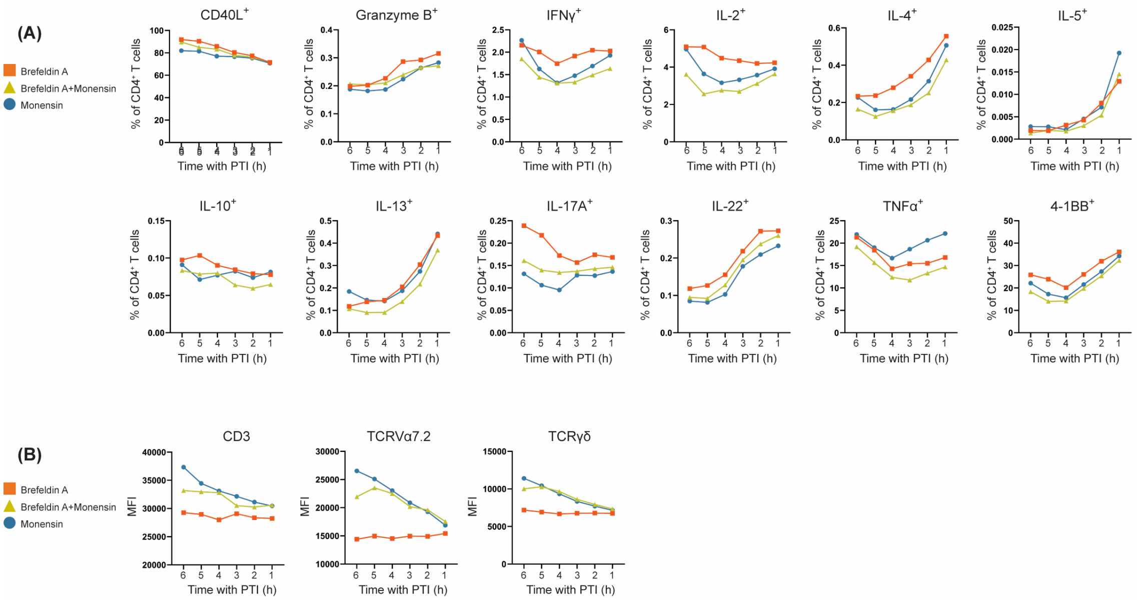


**Online Figure 8: Comparison of protein transport inhibitors (PTIs).** PBMC were stimulated with αCD3/αCD28 for 6h. Brefeldin A and/or BD GolgiStop™ (containing monensin) were added for different durations. (A) Quantification of co-stimulatory markers, granzyme and cytokines as a percentage of $CD4^+$ T cells. (B) Median fluorescence intensity (MFI) of CD3 – Pacific Blue, TCRVα7.2 – BV786 and TCRγδ – PE/Fire 810. Mean data from 2 healthy donors.

**Online Table 1. Instrument configuration** of a 5-laser Cytek® Aurora spectral analyzer (Cytek Biosciences). Fluorochromes used in the final panel are listed next to corresponding detectors.

| Ultraviolet (355 nm - 20 mW) | | | | Violet (405 nm - 100 mW) | | | | Blue (488 nm - 50 mW) | | | | Yellow Green (561 nm - 50 mW) | | | | Red (640 nm - 80 mW) | | | |
|---|---|---|---|---|---|---|---|---|---|---|---|---|---|---|---|---|---|---|---|
| Detector | Center Wavelength (nm) | Bandwidth (nm) | Fluorochrome | Detector | Center Wavelength (nm) | Bandwidth (nm) | Fluorochrome | Detector | Center Wavelength (nm) | Bandwidth (nm) | Fluorochrome | Detector | Center Wavelength (nm) | Bandwidth (nm) | Fluorochrome | Detector | Center Wavelength (nm) | Bandwidth (nm) | Fluorochrome |
| UV1 | 372.5 | 15 | Spark UV 387 | | | | | | | | | | | | | | | | |
| UV2 | 387.5 | 15 | BUV395 | | | | | | | | | | | | | | | | |
| UV3 | 427.5 | 15 | | V1 | 427.5 | 15 | BV421 | | | | | | | | | | | | |
| UV4 | 443 | 15 | | V2 | 443 | 15 | Super Bright 436 | | | | | | | | | | | | |
| UV5 | 458 | 15 | | V3 | 458 | 15 | Pacific Blue | | | | | | | | | | | | |
| UV6 | 473 | 15 | LIVE DEAD Fixable Blue | V4 | 473 | 15 | | | | | | | | | | | | | |
| UV7 | 514 | 28 | BUV496 | V5 | 508 | 20 | BV480 | B1 | 508 | 20 | Spark Blue 515 | | | | | | | | |
| | | | | V6 | 524.5 | 17 | | B2 | 524.5 | 17 | | | | | | | | | |
| UV8 | 542 | 28 | | V7 | 541.5 | 17 | BV510 | B3 | 541.5 | 17 | RB545 | | | | | | | | |
| UV9 | 581.5 | 31 | BUV563 | V8 | 580.5 | 19 | | B4 | 580.5 | 19 | | YG1 | 577 | 20 | PE | | | | |
| | | | | V9 | 598 | 20 | | B5 | 598 | 20 | | YG2 | 598 | 20 | RY586 | | | | |
| UV10 | 612.5 | 31 | BUV615 | V10 | 615 | 20 | BV605 | B6 | 615 | 20 | RB613 | YG3 | 615 | 20 | RY610 | | | | |
| UV11 | 664 | 27 | BUV661 | V11 | 664 | 27 | BV650 | B7 | 661 | 17 | | YG4 | 661 | 17 | | R1 | 661 | 17 | APC |
| | | | | | | | | B8 | 679 | 18 | PerCP | YG5 | 679 | 18 | PE/Cy5 | R2 | 679 | 18 | Alexa Fluor 647 |
| UV12 | 691.5 | 28 | | V12 | 691.5 | 28 | | B9 | 697 | 19 | | YG6 | 697 | 19 | | R3 | 697 | 19 | Spark NIR 685 |
| UV13 | 720 | 29 | | V13 | 720 | 29 | | B10 | 717 | 20 | PerCP/eFluor 710 | YG7 | 720 | 29 | PE/Fire700 | R4 | 717 | 20 | R718 |
| | | | | | | | | B11 | 738 | 21 | | | | | | R5 | 738 | 21 | |
| UV14 | 749.5 | 30 | BUV737 | V14 | 749.5 | 30 | BV750 | B12 | 760 | 23 | RB744 | YG8 | 749.5 | 30 | | R6 | 760 | 23 | |
| UV15 | 779.5 | 30 | | V15 | 779.5 | 30 | BV786 | B13 | 783 | 23 | | YG9 | 779.5 | 30 | PE/Cy7 | R7 | 783 | 23 | |
| UV16 | 811.5 | 34 | BUV805 | V16 | 811.5 | 34 | | B14 | 811.5 | 34 | RB780 | YG10 | 811.5 | 34 | PE/Fire 810 | R8 | 811.5 | 34 | APC/Fire 810 |

### Online Table 2. Reagent information

| Specificity | Clone | Fluorochrome | Vendor | Cat# | Titration (final dilution) |
|---|---|---|---|---|---|
| *Surface staining* | | | | | |
| Dead cells | n/a | LIVE DEAD Blue | Invitrogen | L34961 | 1:500 |
| CD7 | M-T701 | BUV805 | BD | 742002 | 1:80 |
| CD8 | RB545 | SK1 | BD | 756508 | 1:80 |
| CD27 | L128 | BUV496 | BD | 750168 | 1:160 |
| CD38 | HIT2 | APC/Fire 810 | Biolegend | 303550 | 1:40 |
| CD45 | HI30 | PerCP | Biolegend | 982318 | 1:80 |
| CD45RA | HI100 | Spark UV 387 | Biolegend | 304180 | 1:20 |
| CD56 | NCAM16.2 | BUV661 | BD | 750478 | 1:640 |
| CD69 | FN50 | Super Bright 436 | eBioscience | 62-0699-42 | 1:20 |
| CD103 | Ber-ACT8 | PE/Fire 700 | Biolegend | 350240 | 1:80 |
| CD161 | HP-3G10 | BUV563 | BD | 749223 | 1:20 |
| CD197 (CCR7) | G043H7 | Spark NIR 685 | Biolegend | 353258 | 1:20 |
| CD294 (CRTH2) | BM16 | BUV615 | BD | 751216 | 1:40 |
| HLA-DR | G46-6 | BV480 | BD | 566113 | 1:320 |
| *Intracellular staining* | | | | | |
| CD3 | UCHT1 | Pacific Blue | Biolegend | 980004 | 1:200 |
| CD4 | OKT4 | BV510 | Biolegend | 317444 | 1:200 |
| CD137 (4-1BB) | 4B4-1 | BV650 | Biolegend | 309828 | 1:800 |
| CD154 (CD40L) | 2431 | BV605 | Biolegend | 310826 | 1:200 |
| FoxP3 | PCH101 | PE/Cy5 | eBioscience | 15-4776-42 | 1:1600 |
| Granzyme B | GB11 | RB744 | BD | 570498 | 1:800 |
| Helios | 22F6 | PerCP/eFluor 710 | eBioscience | 46-9883-42 | 1:100 |
| IFNγ | B27 | BV750 | BD | 566357 | 1:1600 |
| IL-2 | MQ1-17HI2 | BUV737 | BD | 612836 | 1:800 |
| IL-4 | MP4-25D2 | RB780 | BD | 569087 | 1:800 |
| IL-5 | TRFK5 | RY586 | BD | 568530 | 1:400 |
| IL-9 | MH9A3 | BV421 | BD | 564254 | 1:800 |
| IL-10 | JES3-19F1 | R718 | BD | 567059 | 1:200 |
| IL-13 | JES10-5A2 | APC | Biolegend | 501907 | 1:800 |
| IL-17A | N49-653 | RY610 | BD | 571173 | 1:3200 |
| IL-21 | 3A3-N2.1 | AF647 | BD | 560493 | 1:100 |
| IL-22 | 22URTI | PE/Cy7 | eBioscience | 25-7229-42 | 1:1600 |
| Ki-67 | B56 | BUV395 | BD | 564071 | 1:1600 |
| RORyt | Q21-559 | PE | BD | 563081 | 1:200 |
| T-bet | 4B10 | RB613 | BD | 571286 | 1:1600 |
| TCRγ/δ | B1 | PE/Fire 810 | Biolegend | 331243 | 1:3200 |
| TCR Vα7.2 | OF-5A12 | BV786 | BD | 749488 | 1:800 |
| TNFα | MAb11 | Spark Blue 515 | Biolegend | 502956 | 1:200 |
